# USP18-STAT2 axis enhances hepatic resilience under proteotoxic stress

**DOI:** 10.64898/2026.07.11.737961

**Authors:** Abhishek Sen, Saheli Chowdhury, Partha Chakrabarti

## Abstract

The liver is a metabolic hub with a high protein turnover that renders it uniquely susceptible to proteotoxic stress. Perturbation of proteostasis, either by proteasomal inhibitors or in chronic liver diseases, could adversely impact liver physiology. Here, we show that proteasomal inhibition unexpectedly suppresses basal type I interferon (IFN-I) signaling in the murine liver. Proteasomal inhibition by bortezomib selectively downregulates a subset of interferon-stimulated genes (ISGs), among which USP18 and ISG15 emerge as critical determinants of hepatocellular survival. We identify USP18 as a central cytoprotective factor that prevents proteotoxic apoptosis independently of its deubiquitinase activity, but strictly requires its scaffolding function mediated by isoleucine-60 and interaction with STAT2. Mechanistically, proteotoxic stress disrupts IRF9 nuclear translocation, attenuating USP18 transcription, and drives USP18 and other ISGs into insoluble aggregates with kinetics distinct from canonical IFN-I–induced insolubility. Strikingly, IFN-I priming preserves ISG solubility, restores USP18 abundance, and confers resistance to proteotoxic cell death. Together, these findings uncover an unanticipated link between proteostasis and innate immune signaling, and establish the USP18-STAT2 axis to enhance hepatic resilience under proteotoxic stress.

## INTRODUCTION

The liver orchestrates essential metabolic processes and maintains systemic proteostasis through tightly coordinated control of protein synthesis, folding, degradation, and secretion. Disruption of the latter network contributes to the pathogenesis of major chronic liver diseases, including metabolic dysfunction–associated steatotic liver disease (MASLD), alcoholic liver disease (ALD), viral hepatitis, and hepatocellular carcinoma (HCC) [1,2]. A hallmark of these conditions is the formation of Mallory–Denk bodies (MDBs)—cytoplasmic inclusions enriched in misfolded keratins (KRT8/18), ubiquitin, p62, and molecular chaperones [3]—underscoring proteostasis collapse as a defining pathological feature.

The ubiquitin–proteasome system (UPS) is a key determinant of hepatic protein quality control, responsible for the clearance of damaged or misfolded proteins. Impairment of the UPS promotes MDB formation, increases susceptibility to proteotoxic stress, and contributes to metabolic and inflammatory liver pathology [4–6]. Proteasome dysfunction further induces ER stress and activates the ASK1–JNK/p38 signaling axis while suppressing PPARγ- and Nrf2-dependent antioxidant responses, collectively driving hepatocellular injury and fibrosis. Combined ASK1 inhibition and PPARγ activation can counteract these deleterious effects [7]. In ALD, ethanol-induced oxidative stress compromises proteasomal activity, leading to accumulation of oxidized proteins, while ubiquitin has emerged as a potential biomarker of hepatocellular injury [8–9]. Clinically, the sensitivity of hepatocytes to proteasome inhibition is evident from cases of severe liver injury in multiple myeloma patients treated with bortezomib [10–12]. Consistent with this, the deubiquitinating enzyme JOSD1 has recently been identified as a protective factor against bortezomib-induced hepatic proteotoxicity [13].

The present study aimed to identify and characterize previously unrecognized pathways that modulate proteotoxic stress in hepatocytes. We report that the basal type I interferon (IFN-I) signaling pathway is unexpectedly downregulated upon proteasomal inhibition, despite its canonical role in cellular defense. Restoring IFN-I signaling markedly reduces hepatocellular proteotoxicity, and among the ISGs, the deubiquitinase and deISGylase USP18 emerges as a central molecular determinant of this cytoprotective response.

## RESULTS

### Proteasomal Inhibition Attenuates Hepatic Type I Interferon Signalling

To elucidate the adverse consequences of proteasomal inhibition, C57BL/6 mice were administered with the proteasome inhibitor bortezomib via intraperitoneal injection. Bortezomib caused a significant liver injury compared to other internal organs, such as the lungs and kidneys (Fig S1A). We next conducted a transcriptomic analysis of hepatic tissues of these mice^7^. As expected, gene ontology enrichment revealed upregulation of biological processes associated with proteasomal stress and protein quality control. Unexpectedly, however, we observed significant downregulation of the IFN-I signalling pathway (Figure S1B). This suppression was characterized by a marked reduction in the expressions of multiple interferon-stimulated genes (ISGs) (Figures 1A, S1C). To validate these transcriptomic findings, mice were treated with bortezomib for varying durations up to 16 hours. Quantitative analyses demonstrated a time-dependent decrease in the mRNA expression of selected genes involved in IFN-I signalling—USP18, STAT1, STAT2, IRF9, RTP4, and ISG15—with the exception of ISG15, which remained largely unaltered (Figure 1B). Consistent with these results, the protein levels of USP18, STAT1, and STAT2 were also progressively reduced following bortezomib treatment (Figure 1C). Proteasomal inhibitor MG132 also elicited a similar response in downregulating ISGs in the IFN-I signalling pathway in cell autonomous manner (Figure S1D). These findings collectively suggest that proteasomal inhibition suppresses basal expression of a subset of ISGs involved in the IFN-I signalling pathway in hepatocytes. Given that ISG transcription in the context of the type 1 IFN pathway is primarily regulated by the transcription factor IRF9, which forms the ISGF3 complex with STAT1 and STAT2, or a heterodimeric complex with STAT2 [14–15], we next examined whether proteasomal blockade perturbs IRF9 dynamics in hepatocytes. In the human hepatocellular carcinoma cell line HepG2, treatment with MG132 led to an impairment of IRF9 nuclear localization. Notably, analysis of nuclear extracts confirmed a substantial reduction in nuclear IRF9 levels upon proteasomal inhibition (Figure 1D). It also abolished IFN-induced nuclear translocation of IRF9 (Figure 1E). To further assess whether proteasomal inhibition-mediated ISG suppression is dependent on IRF9, we performed siRNA-mediated knockdown of IRF9 in HepG2 cells with or without MG132 treatment. Silencing IRF9 significantly decreased ISG transcript levels, and subsequent proteasomal blockade did not further suppress ISG expression, indicating an IRF9-dependent mechanism (Figure 1F). ISG15 expression, however, remained largely unaffected at the transcript level. In contrast, protein levels of ISGs were markedly reduced following MG132 treatment even in IRF9-depleted cells (Figure 1G). Collectively, these data indicate that proteasomal inhibition constrains basal IFN-I signalling by preventing IRF9 nuclear localization and thereby repressing ISG transcription. Furthermore, the persistence of ISG protein loss in the absence of IRF9 suggests an additional post-transcriptional mechanism contributing to the proteasome-dependent downregulation of interferon-stimulated proteins. Consistently, proteasomal inhibition in combination with global transcriptional inhibitor Actinomycin D resulted in a greater loss of ISG proteins compared with Actinomycin D treatment alone (Figure 1H). Moreover, concomitant treatment with MG132 and Actinomycin D did not elicit any significant alteration in the ISG transcript patterns relative to the only Actinomycin D treatment group (Figure 1I).

**Figure 1.**
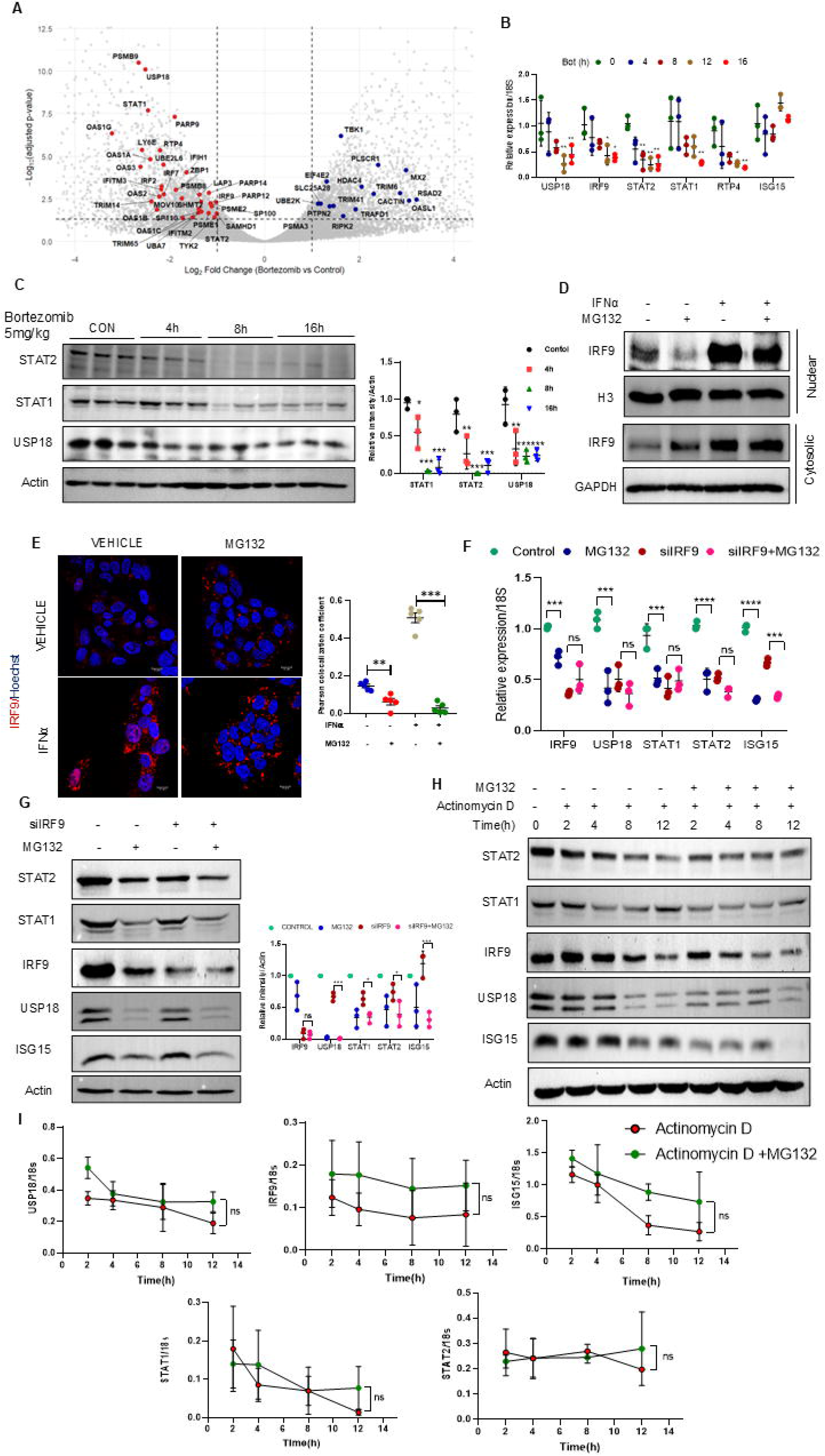
Proteasome blockade attenuates the hepatic IFN-I signaling. (A) Volcano plots of ISGs showing log₂ fold change versus −log₁₀ (adjusted p-value) on proteasomal inhibition by 5 mg/Kg bortezomib treatment in C57BL/6 mice. Genes with adjusted p-value < 0.05, and |log₂ fold change| ≥ 1 were considered significant. (B, C) Gene expression (B), and Western blot (C) of ISGs from liver tissue of C57BL/6 mice intraperitoneally injected with 5 mg/kg dose of bortezomib for 4h, 8h and 16 h (n=3 mice/group). Densitometric analysis of relative intensity normalized by actin given right. (D) Nuclear and cytosolic fractions from HepG2 cells treated with 5μM of MG132 for 4h, followed by 500 IU/ml of IFNα treatment for 1h. (E) Nuclear localization of IRF9 in HepG2 cells treated with 5μM of MG132 for 4 h, followed by 500 IU/ml of IFNα treatment for 1h. (F-G) Relative gene expression (F) and Western blot (G) analysis of ISGs under IRF9 knockdown conditions, treated with 5μM doses of MG132 for 16h. Densitometric analysis of relative intensity of ISGs normalized with actin is given on the right (For 3 independent experiments). (H, I) HepG2 cells were treated with 5µg/ml of Actinomycin D and with 5µM MG132 for the indicated time periods. Protein levels were measured by Western blot (H), and relative transcript levels were measured by qPCR analysis (I). Values represent mean ± SD. Statistical significance: **P < 0.01; ***P < 0.001, *P < 0.05; NS, non-significant comparisons. Scale bar = 10 µm.

### ISGs Aggregates distribute to the Insoluble Fraction under Proteotoxic Stress

To examine the molecular mechanisms underlying post-transcriptional regulation of ISGs, we investigated the potential involvement of protein stability and degradation pathways. Given that proteasomal blockade typically induces compensatory activation of autophagy [16–17], we pharmacologically inhibited autophagy using bafilomycin A. Although treatment with MG132 led to autophagic activation, co-treatment with bafilomycin A failed to rescue proteasomal inhibition-induced suppression of ISG protein levels (Figure S1E). Because proteasomal inhibition is known to elicit apoptosis^7^, we next examined the contribution of apoptosis to ISG regulation. Inhibition of caspase activity using the pan-caspase inhibitor Z-VAD-FMK efficiently prevented cell death but did not restore ISG protein expressions (Figure S1F). The lack of rescue upon inhibition of autophagy and apoptosis, prompted us to explore the possibility of ISG aggregation, as proteasomal inhibition often triggers the unfolded protein response (UPR) and the formation of perinuclear aggresomes mediated by the autophagy receptor p62 [18]. To determine the temporal dynamics of ISG expression under proteotoxic conditions, HepG2 cells were treated with MG132 for up to 16 hours. As expected, transcript levels of representative ISGs—including USP18, STAT1, STAT2, IRF9, and ISG15—showed a progressive decline, whereas p62, a classical aggresomal marker, remained relatively unaltered (Figure 2A). Notably, with prolonged proteasomal inhibition, endogenous ISG proteins gradually shifted from the detergent-soluble to the detergent-insoluble fraction, mirroring the kinetics of p62 redistribution (Figure 2B). These observations suggest that proteasomal inhibition suppresses ISGs through dual mechanisms: transcriptional repression via the IRF9 regulatory axis, and post-transcriptional aggregation leading to insolubility of ISG proteins.

**Figure 2.**
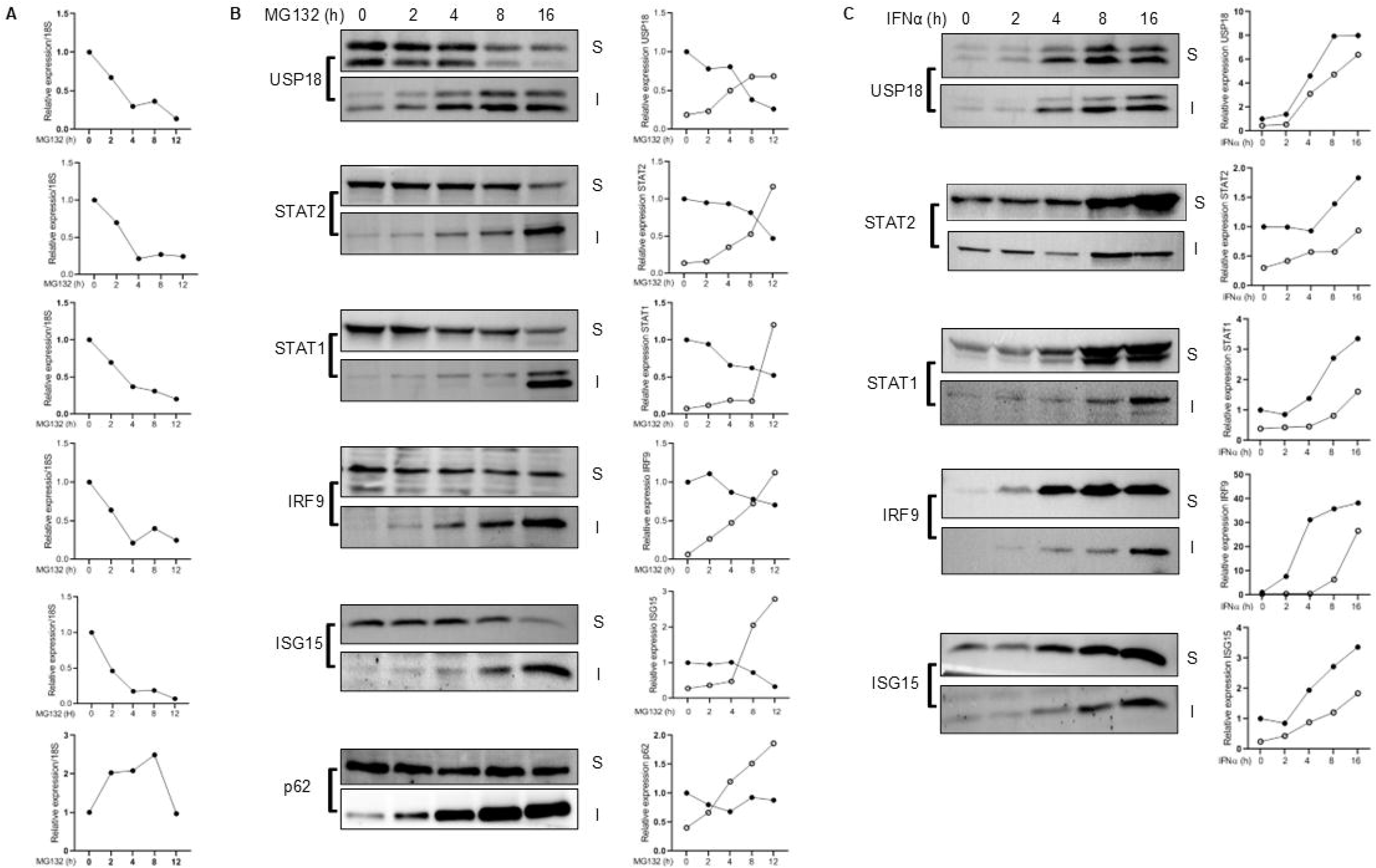
Partitioning of ISGs into insoluble fractions by proteasomal inhibition and IFNα stimulation. (A, B) HepG2 cells were treated with 5µM of MG132 for the indicated time periods, and total cellular RNA, soluble and insoluble fractions of proteins were isolated. Gene expressions of indicated ISGs and p62 were determined by qPCR (A). Left panels: The soluble and insoluble fractions were subjected to Western blot analysis. Right panels: Relative abundance of soluble and insoluble proteins was determined by densitometric analysis, considering the soluble fraction at the basal state as 1 (B). (C) Left panels: HepG2 cells were treated with 500 IU/ml of IFNα for 2h, 4h, 8h, and 16h, and the detergent-soluble and detergent-insoluble fractions were subjected to Western blot analysis. Right panels: Relative abundance of soluble and insoluble proteins was determined by densitometric analysis, considering the soluble fraction at the basal state as 1. S- Detergent soluble fraction; I- Detergent insoluble fraction.

We next asked whether interferon-induced ISG expression could similarly promote protein insolubility. Interestingly, stimulation with IFNα resulted in a time-dependent accumulation of ISGs in both soluble and detergent insoluble fractions (Figure 2C). Thus, IFNα-mediated expression appears to drive a fraction of ISG proteins toward the insoluble pool, possibly due to concentration-dependent mass action effects. Importantly, the kinetics of insolubility differed between proteotoxic stress and IFNα stimulation: under proteasomal inhibition, ISGs were depleted from the soluble pool and enriched in the detergent insoluble fraction, whereas IFNα treatment increased ISG abundance in both compartments. We next examined whether ISGs have any analogous structural motifs that make them susceptible to aggregation. Since their full structures are not publicly available, we used the Alphafold platform to delineate the tertiary protein structures. Interestingly, we saw nearly all ISGs (USP18, STAT1, STAT2, IRF9, and ISG15) have a long stretch of intrinsically disordered region (IDR) like classical aggresome marker p62/SQSTM1 (Figure S2), suggesting that IDR’s in these ISGs might contribute to their propensity to form aggresome.

### ISG USP18 is the Major Contributor in Mitigating Proteotoxic Death

Proteasomal inhibition disrupts cellular proteostasis, resulting in the accumulation of misfolded damaged proteins, ultimately triggering hepatocellular injury [7]. The suppression of basal IFN-I response following proteasomal blockade prompted us to investigate how ISGs influence hepatic proteotoxicity. To address this, we individually silenced key components of the IFN signaling pathway—STAT1, STAT2, IRF9, USP18, and ISG15, and cells were then subjected to MG132 treatment to induce proteotoxic apoptosis. Among the candidates tested, knockdown of USP18 and ISG15 markedly exacerbated the expression of cleaved caspase-3 (CC3), a hallmark of apoptosis (Figures 3A–E). Consistently, quantitative assays assessing apoptosis and cell viability further confirmed the protective roles of USP18 and ISG15 during proteotoxic stress (Figure 3F). USP18 functions as both a deubiquitinase and deISGylase, and serves as a critical negative regulator of IFN signaling [19–20], whereas ISG15 is a canonical effector ISG mediating IFN-induced response [21]. Our data, therefore, suggest that both USP18 and ISG15 play essential roles in mitigating proteotoxicity. To delineate whether USP18 and ISG15 expressions had confounding effects on other ISGs, we either overexpressed or knocked down USP18 and ISG15. Ectopic expression or silencing of either of them had no significant impact on the expression of other ISGs (Figures S3A, B). To further demarcate their relative contributions in proteotoxicity, we performed dual transfection experiments in which USP18 was overexpressed under conditions of ISG15 depletion, and vice versa. Ectopic expression of USP18 effectively suppressed CC3 accumulation even in the absence of ISG15 (Figure 3G). In contrast, overexpression of ISG15 failed to rescue proteotoxic apoptosis when USP18 was silenced (Figure 3H). Of note, overexpression of ISG15 alone could suppress proteotoxicity (Figure S3C). Microscopic analyses of cleaved caspase-3 (CC3) immunostaining corroborated these findings (Figure 3I). Collectively, these results establish that USP18 is both necessary and sufficient to mitigate proteotoxic cell death, underscoring its dominant protective role among ISGs.

**Figure 3.**
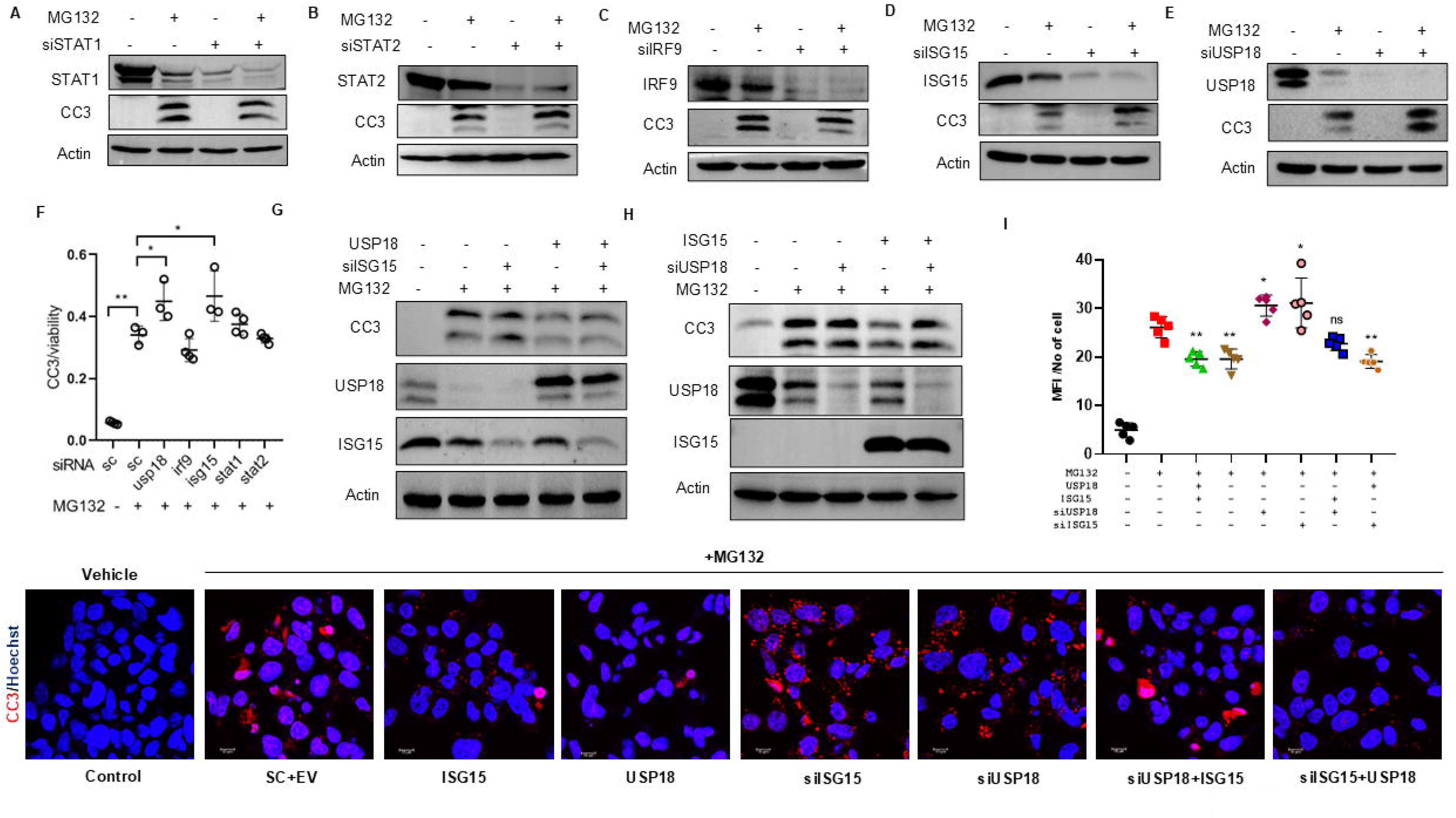
ISG USP18 mitigates proteotoxicity. (A-E) Western blot analysis of HepG2 cell lysates transfected with siSTAT1 (A), siSTAT2 (B), siIRF9 (C), siUSP18 (D), and siISG15 (E), followed by 5µM of MG132 treatment for 16h. (F) HepG2 cells were transfected with siSTAT1, siSTAT2, siIRF9, siUSP18, and siISG15 in 96 well plate, followed by 5μM of MG132 treatment for 16h. The ApoLive-Glo™ multiplex assay was performed in triplicate to simultaneously measure apoptosis and viability. Representative data shown for three independent experiments. (G) Western blot analysis of HepG2 cells lysate with USP18 overexpression, and ISG15 knockdown along with 16 h of 5 μM of MG132 treatment. (H) Western blot analysis of HepG2 cells lysate co-transfected with siUSP18 and ISG15 for 48h, followed by 16h of 5μM of MG132 treatment. (I) Lower panel: HepG2 cells transfected with siRNA and plasmid constructs as indicated, followed by 16h of 5μM of MG132 treatment. Cells were then stained for CC3, and nuclei were stained with Hoechst. Upper panel: Mean fluorescence intensity (MFI) of CC3 was determined (5 fields/group). Statistical comparison was done with respect to MG132-treated cells. sc- Scrambled siRNA; EV- empty vector. Values represent mean ± SD. Statistical significance: **p< 0.01; *p< 0.05. Scale bar = 10 µm.

### USP18 forms distinct, reversible intracellular aggregates under proteotoxic stress

ISGs partition into the detergent insoluble protein fraction following proteasomal blockade. To determine whether these insoluble ISGs form stable aggregates, cells were treated with MG132 for varying durations, followed by withdrawal of the media to relieve proteotoxic stress. Interestingly, the insolubility of ISGs decreased upon removal of proteotoxic stress, and the extent of this reduction was inversely correlated with the duration of proteotoxic stress (Figure 4A), indicating that ISG insolubility is a dynamic and reversible process. Given the protective role of USP18 during proteotoxic stress and its intrinsic tendency to form aggregates, we observed that USP18 aggregates formed during proteasomal inhibition were disrupted by treatment with 5% 1,6-hexanediol (1,6-HD), further indicating the reversibility of the aggregates (Figure 4B). We next ectopically expressed USP18 in HepG2 cells, overexpression of USP18 led to its accumulation in the detergent insoluble fraction, with a substantial fraction in the soluble pool, even following MG132 treatment (Figure 4C). Immunofluorescence analysis revealed that USP18 accumulated as discrete cytoplasmic puncta in a dose-dependent manner, upon MG132 treatment, indicative of phase-separated protein condensates (Figure 4D). To examine whether USP18 aggregates overlap with p62-containing aggresomes, cells were co-stained for both proteins. Whereas p62 formed characteristic perinuclear aggresomes following proteasomal inhibition, USP18 puncta remained diffusely distributed throughout the cytoplasm without co-localization with p62 (Figure 4E). To further evaluate USP18 aggregation propensity, we performed an in vitro aggregation assay using immunoprecipitated flag-tagged USP18 [22]. Purified USP18 protein spontaneously formed insoluble protein droplets, which were markedly enhanced when USP18 was isolated from MG132-treated cells (Figure 4F). Collectively, these observations demonstrate that under proteotoxic stress, USP18 forms reversible, phase-separated protein droplets that are distinct from p62 aggresomes.

**Figure 4.**
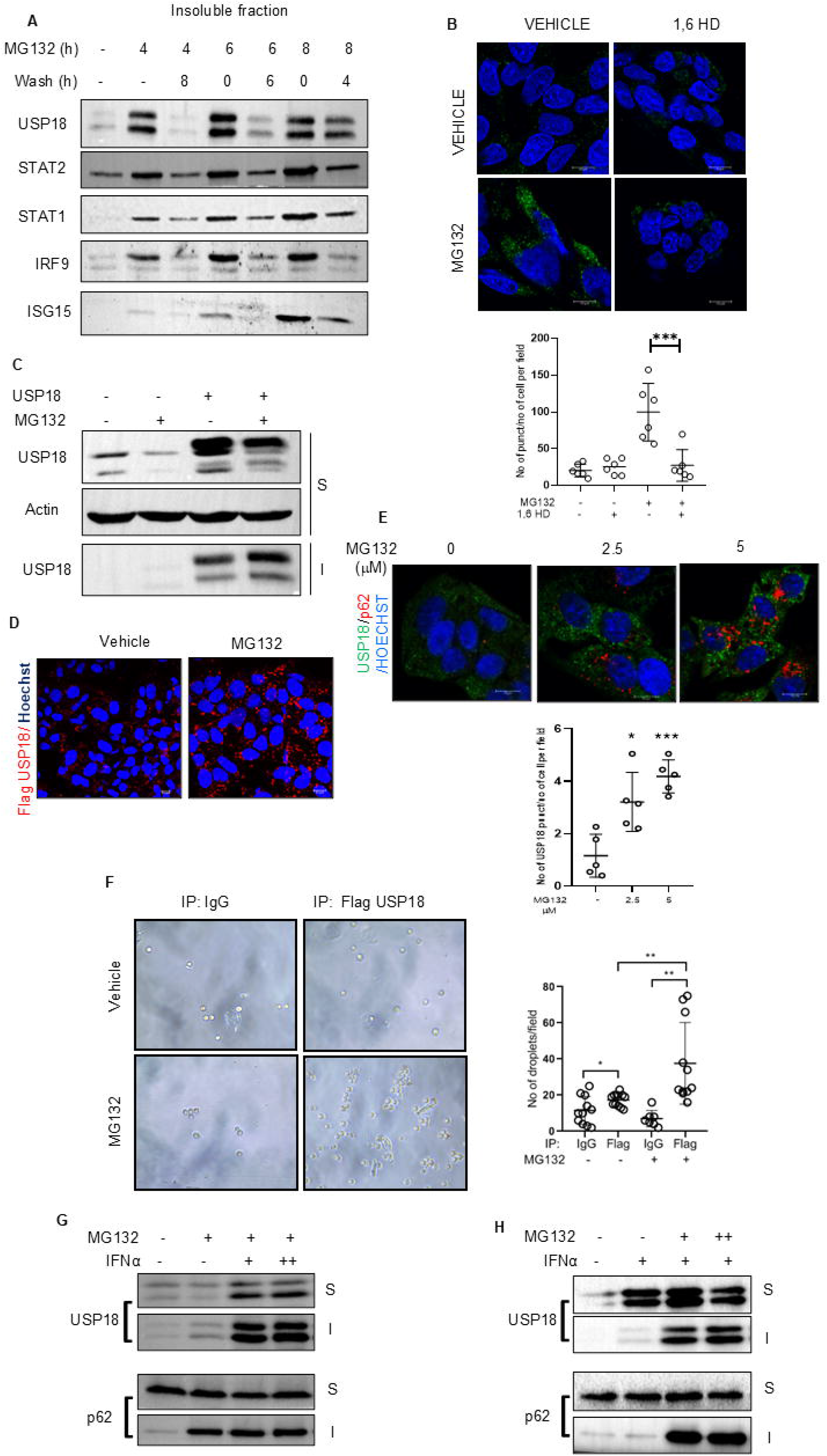
Proteotoxicity induces distinct protein aggregates of USP18. (A) HepG2 cells were treated with 5μM of MG132 for 4h, 6h, and 8h, followed by washing the cells for 8h, 6h and 4h, respectively. Expressions of ISGs in the insoluble fractions were analyzed by Western blotting. (B) HepG2 cells were treated with 5µM MG132 for 6h followed by treatment with 5% 1,6-HD for 2 min. USP18 was stained, and nuclei were stained with Hoechst, visualized by confocal microscopy. Statistical analysis of no of puncta of USP18/ no of cells for 6 fields/group given at the bottom. (C) USP18 was overexpressed in HepG2 cells and treated with 5µM of MG132 for 8h. Levels of USP18 in the soluble and insoluble fractions were determined by Western blotting. (D) HepG2 cells were transfected with flag-tagged USP18, followed by 8h of MG132 treatment. USP18 was stained with anti-flag antibody and visualized by confocal microscopy. (E) HepG2 cells were treated with 2.5 μM and 5 μM of MG132 for 8h. Cells were stained for USP18 and p62. Statistical analysis of no of USP18 puncta/no of cells for 5 fields/group is given in the bottom. The comparison was done with respect to control (F) Left panel: HepG2 cells were transfected with flag-tagged USP18 and treated with 5μM MG132 for 12h. USP18 was immunoprecipitated using anti-flag tag antibody and incubated for 1h at 37^0^C. Aggregate formation was visualized by bright-field microscopy. Right panel: Number of aggregates per microscopic field counted (8 fields/group). (G-H) HepG2 cells were pretreated with 250 IU/ml and 500 IU/ml of IFNα for 4h, followed by 12h of 5µM MG132 treatment (G). Cells were pretreated with 500IU/ml of IFNα for 4h, followed by 12h of 5µM and 10µM MG132 treatment (H). Levels of USP18 and p62 in the soluble and insoluble fractions were determined by Western blotting. S-detergent soluble fraction; I- detergent insoluble fraction. Values represent mean ± SD. Statistical significance: **p< 0.01; *p < 0.05; Scale bar = 10 µm.

We next investigated whether IFN-I signaling modulates USP18 solubility under proteotoxic conditions. Cells were treated with increasing concentrations of either IFNα or MG132 while keeping the other constant. IFNα treatment did not fully prevent USP18 insolubility during proteasomal inhibition but significantly increased the proportion of USP18 present in the soluble fraction (Figures 4G, H). Notably, USP18 remained predominantly soluble under IFNα treatment alone (Figure 4H). In contrast, the aggregation of p62 was unaffected by IFNα. These results indicate that the availability of fraction of soluble USP18 under IFN-I stimulation might be critical for its cytoprotective function despite partial presence of insoluble USP18, whereas nearly complete loss of soluble USP18 upon gene silencing exacerbates proteotoxic cell death.

### IFN-I Mitigates Proteotoxicity via USP18

Given that proteotoxic stress suppresses basal IFN-I responses, which are largely regulated by the ISG USP18, we investigated whether IFN-I stimulation could attenuate proteotoxicity in a USP18-dependent manner. To this end, HepG2 cells were pretreated with increasing concentrations of IFNα followed by a fixed dose of the proteasome inhibitor MG132. IFNα pretreatment conferred marked protection against proteotoxic cell death (Figure 5A). Notably, proteasomal inhibition did not alter canonical IFN-I signaling, as indicated by unchanged phosphorylation of STAT1 and STAT2 (Figure 5B). Consistent with this, ISGylation of cellular proteins remained largely unaffected by MG132, whereas IFNα treatment robustly induced ISGylation (Figure S3D). Since IFNα is known to modulate immunoproteasome function [23], we investigated whether IFNα treatment could enhance proteasomal activity under conditions of proteasomal inhibition. However, dose-dependent IFNα treatment did not improve proteasomal function in the proteasome-inhibited state, as evidenced by the absence of any significant change in total cellular ubiquitination patterns (Figure S3E). To determine whether USP18 is required for the cytoprotective effect of IFNα, USP18 expression was silenced in HepG2 cells. Knockdown of USP18 abolished the protective effect of IFNα against proteotoxicity (Figure 5C). The USP18-dependent protective function of IFNα was further validated microscopically using CC3 immunostaining and live–dead assays. As shown in Figure 5D, depletion of USP18 not only abrogated the protective effect of IFNα but also exacerbated cell death under proteotoxic stress. Consistent with the findings that IFNα induces IRF9 expression and nuclear localization (Figure 1D, E), our results indicate that the cytoprotective effect against proteotoxic stress predominantly depends on USP18 levels. We next evaluated the impact of IFNα on hepatic proteotoxicity in vivo. Mice were administered IFNα for four consecutive days and subsequently challenged with bortezomib for 16 hours (Figure 5E). IFNα-induced expression of USP18 and ISG15 remained unaffected by bortezomib, indicating a sustained IFN response (Figures 5F, G). Furthermore, IFNα pretreatment markedly reduced bortezomib-induced CC3 accumulation in the mouse liver (Figure 5H). Collectively, these findings demonstrate that IFN-I alleviates proteotoxic stress through a USP18-dependent mechanism.

**Figure 5.**
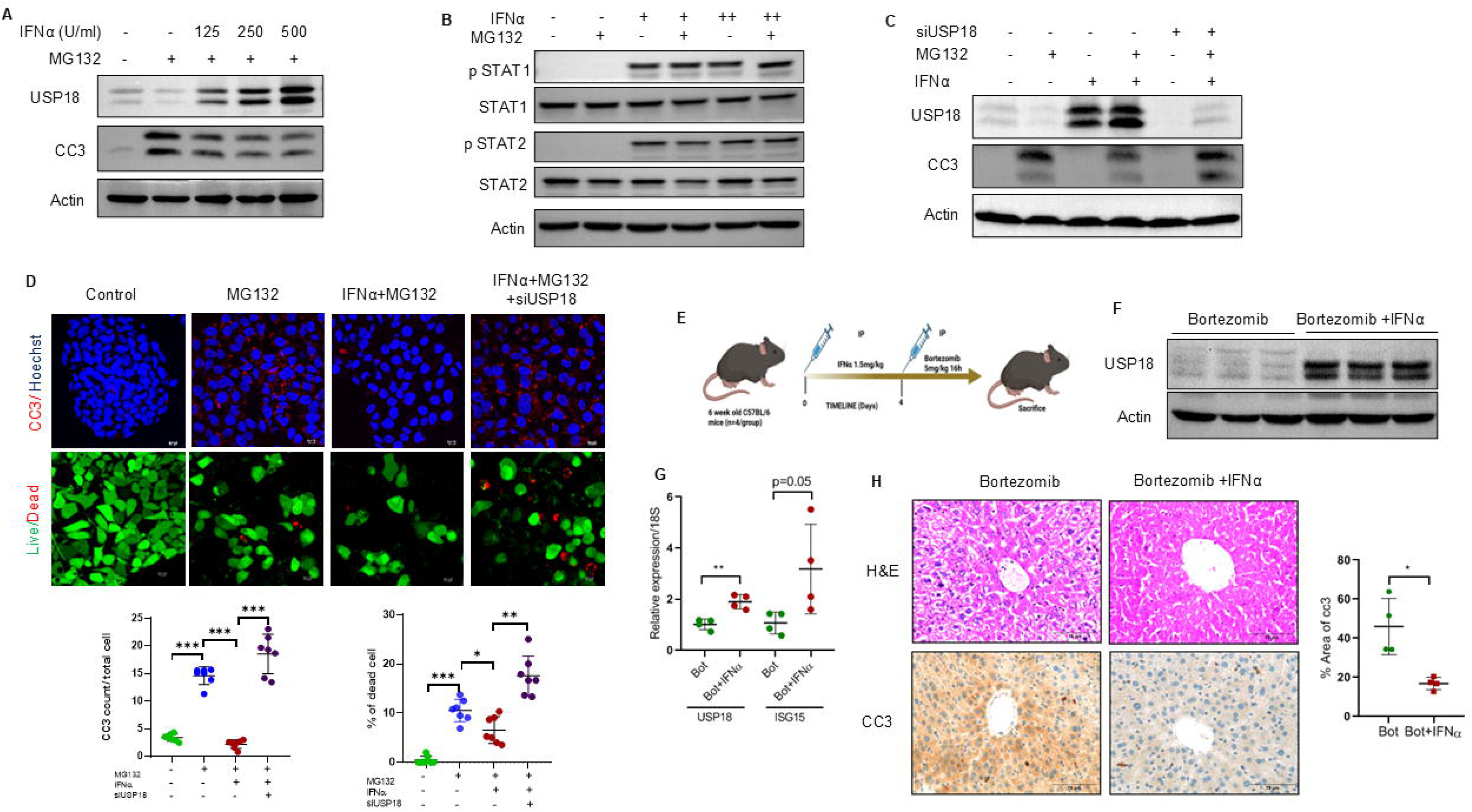
IFN protects from proteotoxic injury via USP18. (A) HepG2 cells were treated with 125, 250, and 500 IU/ml of IFNα for 4h, followed by 5µM of MG132 treatment for 12h. Immunoblotting of USP18 and CC3 was done in the whole cell lysates. (B) HepG2 cells were treated with 5µM of MG132 for 4h, followed by 500 IU/ml of IFNα treatment for 1h and indicated protein levels were determined by Western blotting. (C) Levels of USP18 and CC3 in cells with siRNA-mediated USP18 knockdown, followed by 500 IU/ml of IFNα treatment for 4h and further 12h of 5µM of MG132 treatment. (D) Top panels: Immunostaining of CC3 and live-dead assay in HepG2 cells, with siRNA-mediated USP18 knockdown followed by 4h of 500 IU/ml of IFNα treatment, further followed by 12h of 5µM of MG132 treatment. Scale bar = 10 µm. Bottom panels: The number of CC3 positive cells per field and the percentage of dead cells were determined (7 fields/group). (E) Schematic showing IFNαand bortezomib treatment protocol. Four-week-old male C57BL/6 mice were intraperitoneally injected with 1.5 μg/kg bodyweight of IFNα for 4 days, followed by 16h of 5mg/kg bortezomib. The other group only received bortezomib treatment (n=4 mice/group). (F) Western blot analysis of liver lysate for USP18. (G) qPCR analysis for the expression of ISGs. (H) Haematoxylin and eosin staining (H&E), and IHC for CC3 in bortezomib and bortezomib+ IFNα treated groups. Scale bar=75µm. Right panel: Percentage of CC3 positive area per field was determined (7 fields/mice, n=4 mice/group). Values represent mean ± SD. Statistical significance: **p< 0.01; *p< 0.05.

### USP18 Protects Against Proteasomal Inhibition–mediated Hepatocellular Death

Given our finding that USP18 plays a major role in mitigating proteotoxic stress, we depleted USP18 and challenged HepG2 cells with proteotoxic stress. As shown in Fig S4A, B, knockdown of USP18 augmented proteotoxic death. We next examined whether its cytoprotective expression is broadly modulated across diverse cellular stress responses. HepG2 cells were treated with palmitic acid, doxorubicin, or tunicamycin to induce lipotoxic, genotoxic, and ER stress, respectively. Whereas proteotoxic stress robustly reduced USP18 levels, none of the other stressors altered USP18 abundance (Figure 4C-E), indicating that USP18 regulation is specific to proteotoxicity. Because USP18 overexpression substantially increased the soluble pool of USP18 (Figure 4B), and given that the soluble fraction appears to confer cytoprotection, we assessed whether USP18 alone is sufficient to protect cells from proteotoxicity. HepG2 cells ectopically expressing USP18 did not affect the expressions of other ISGs (Figure 6A). Increasing levels of USP18 expression suppressed proteotoxic stress, as evidenced by attenuation of CC3 levels (Figure 6B). USP18 overexpression similarly diminished cellular caspase-3/7activity (Figure 6C). To further validate the protective role of USP18 in vivo, we delivered USP18 to the mouse liver using recombinant adenovirus-mediated gene transfer, and challenged the animals with bortezomib (Figure 6D). Hepatic USP18 overexpression did not significantly alter the expression of other ISGs, apart from a modest increase in ISG15 (Figure 6E), but it markedly reduced liver injury, as shown by decreased CC3, cleaved PARP, and reduced levels of serum liver injury markers like alanine aminotransferase (ALT) and aspartate aminotransferase (AST) (Figure 6F, G). Immunohistochemistry of liver sections confirmed reduced CC3 staining intensity (Figure 6H). Notably, compared to untreated control mice, USP18 expression was moderately induced in livers receiving control adenovirus carrying eGFP, likely due to virus-mediated IFN-I production. USP18 levels were substantially higher upon USP18 overexpression.

**Figure 6.**
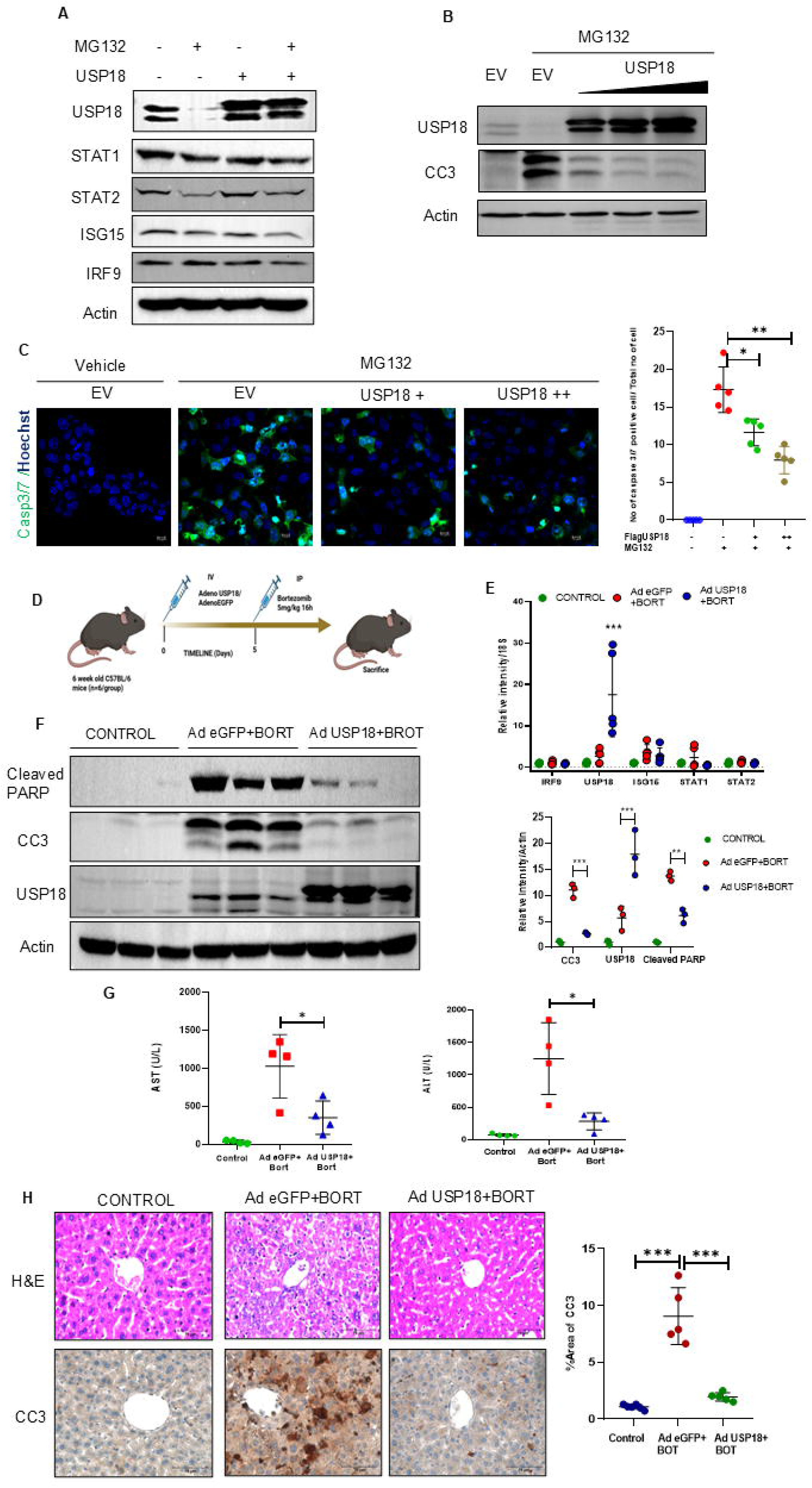
Overexpression of USP18 protects from hepatic proteotoxic death. (A) Western blot analysis of ISGs in HepG2 cells overexpressing USP18 followed by 8h of MG132. (B) USP18 was expressed in escalating dosage in HepG2 cells for 48h, followed by 16h of 5µM of MG132 treatment. Whole cell lysates were analyzed by immunoblotting. (C) Left panel: Caspase3/7 activity in HepG2 cells overexpressed with USP18, followed by 16h of 5µM of MG132 treatment, Scale bar=10µm. Right panel: The number of Caspase3/7 positive cells was counted (5 fields/group). (D) Schematic representation of adenovirus-mediated USP18 overexpression by tail vein injection in C57BL/6 mice. Mice were divided into 3 groups (n=5 mice/group) and injected with recombinant adenovirus expressing eGFP (Ad eGFP, ∼ 2X10^11^ pfu) as a control virus and adenovirus expressing USP18 (Ad USP18, ∼ 2X10^11^ pfu). 5 days after injection, mice were challenged with 5mg/kg body weight of bortezomib via IP injection for 16 h. The control group received a vehicle injection. (E) qPCR analysis of gene expression of ISGs from liver tissues. (F) Western blot analysis of liver tissues. Densitometric analysis of CC3, USP18 and cleaved PARP, normalized to actin (n=3/group). (G) Serum AST and ALT levels from three groups. (H) Haematoxylin and eosin staining (H&E), and IHC analysis to stain CC3 of liver sections, Scale bar=75 µm. Percentage of CC3-positive area was determined (5 fields/mouse, n=5 mice/group). Values represent mean ± SD. Statistical significance: **p< 0.01; ***p< 0.001, *p < 0.05.

### Scaffolding Function of USP18 is Required for the Hepatoprotective Effects Under Proteotoxicity

We next investigated the molecular basis of the cytoprotective function of USP18 during proteotoxic stress. Because USP18 is a deubiquitinating enzyme, we first examined whether its catalytic activity contributes to this effect. Catalytically inactive mutants in which the active-site cysteines were substituted with alanine (C64A) [24] suppressed proteotoxicity to the same extent as wild-type (WT) USP18, demonstrating that enzymatic activity is dispensable for cytoprotection (Figure 7A). These mutants also partitioned into detergent soluble and detergent insoluble fractions similarly to the WT protein (Figure S5A). We next asked whether proteasomal inhibition promotes hyperubiquitination of USP18, thereby driving its insolubility. Proteotoxic stress markedly increased polyubiquitination of endogenous USP18 (Figure 7B). Bioinformatic analysis of the 23 lysine residues using UbPred [25] identified K29, K30, and K37 as the most probable ubiquitination sites. Structural mapping showed that these residues lie within a disordered loop region of the protein (Figure 7C), consistent with the preferential localization of ubiquitination sites in flexible loops [25–27]. Among the corresponding mutants (K29R, K30R, K37R), K37R exhibited reduced ubiquitination relative to WT and the other mutants, identifying K37 as a major ubiquitination site (Figure 7D). However, all three lysine mutants redistributed to the insoluble fraction upon MG132 treatment, indicating that neither ubiquitination nor the lysine residues themselves could determine USP18 insolubility. Consistently, each lysine mutant suppressed proteotoxicity to a level comparable with WT USP18 (Figure S5B).

**Figure 7.**
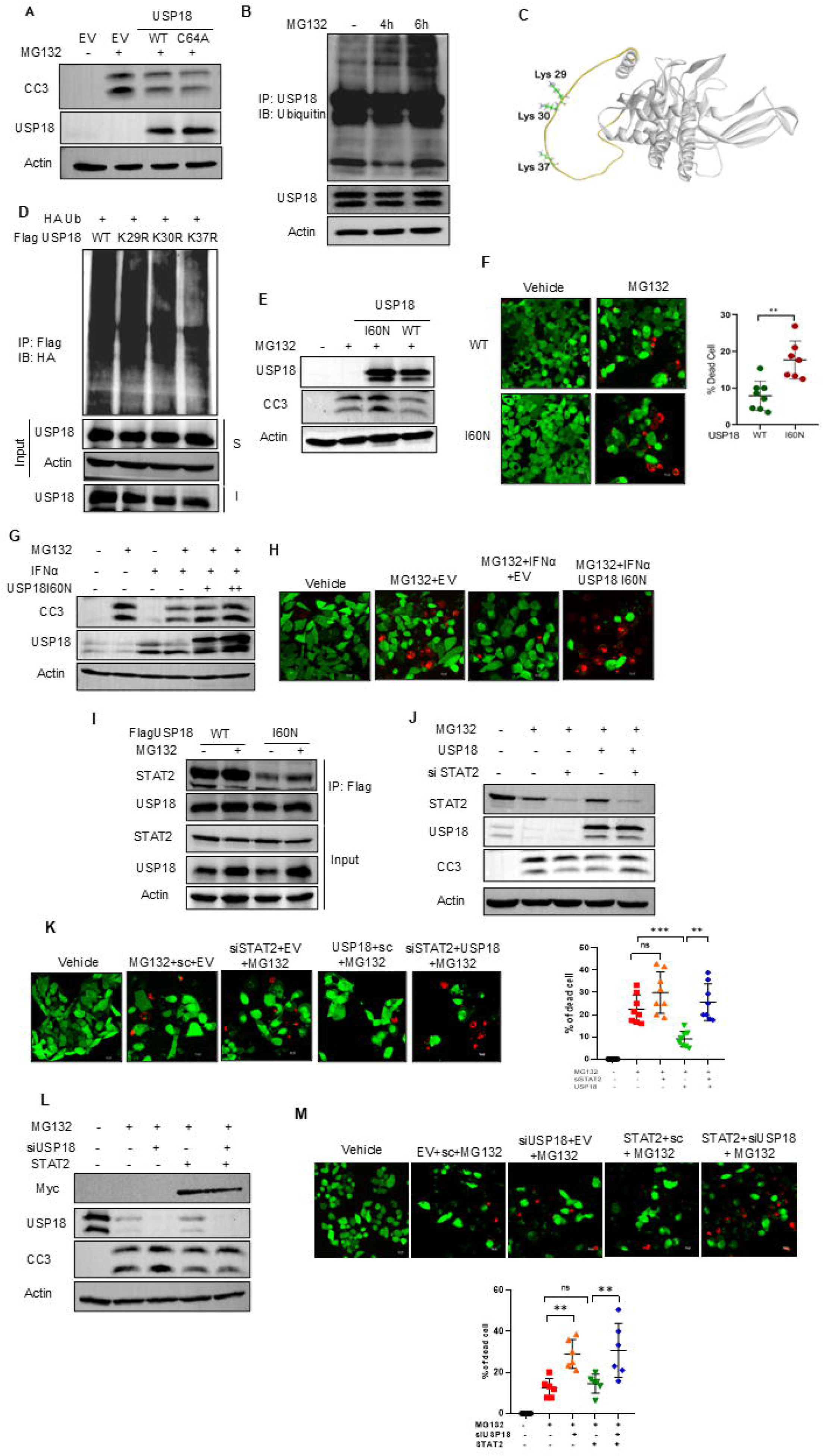
Role of USP18 I60N mutant in proteotoxicity. (A) HepG2 cells were transfected with wild-type and C64A mutant of USP18 and treated with 5µM of MG132 for 16h. Whole cell lysates were analyzed by immunoblotting. (B) HepG2 cells were treated with 5µM of MG132 for 4h and 6h. Immunoprecipitated (IP) using anti- USP18 antibody and immunoblotted (IB) using anti-Ubiquitin antibody. (C) Structure of human USP18 obtained from Alphafold platform showing three predicted lysine residues for ubiquitination. (D) HepG2 cells were transfected with WT Flag tagged USP18 and three lysine mutants, along with HA-ubiquitin (Ub) plasmids and treated with 5µM MG132 for 6h. Immunoprecipitation (IP) was done using anti-Flag tag antibody and immunoblotted (IB) with anti-HA tag antibody. (E-F) Cells were transfected with I60N mutant and treated with 5µM MG132 for 16h. Western blot analysis (E) and live-dead assay (F). The percentage of dead cells was counted for live-dead assay (8 fields/group). (G-H) I60N mutant expressing cells were treated with 500IU/ml IFNα treatment for 4h, followed by MG132 treatment for 12h. Cell lysates were analyzed by Western blot (G), and assayed for live-dead assay (H). The percentage of dead cells was quantified by live-dead assay (8 fields/group) and is shown on the right. (I) HepG2 cells were transfected with Flag tagged USP18 and treated with 5 µM MG132 for 6 h. Immunoprecipitation (IP) was done using anti-Flag tag antibody, and immunoblotted (IB) with anti-STAT2 antibody and USP18 antibody. (J, K) Cells were transfected with siSTAT2 and WT USP18 and treated with 5µM MG132 for 16h. Western blot analysis (J) and live-dead assay (K) were performed. Percentage of dead cells was counted for live-dead assay (8 fields/group) shown at right. (L, M) Cells were transfected with siUSP18 and myc-STAT2 plasmid, and treated with 5µM MG132 for 16h. Western blot analysis (L), and live-dead assay (M) were performed. Percentage of dead cells was counted for live-dead assay (6 fields/group) shown at the bottom right. S-detergent soluble fraction; I-detregent insoluble fraction. sc- Scrambled siRNA; EV- empty vector. Values represent mean ± SD. Statistical significance: **p< 0.01; ***p< 0.001, *p < 0.05; ns denotes non-significant comparisons. Scale bar = 10 µm.

Given that the catalytic and ubiquitin-dependent functions were dispensable, we examined whether USP18’s adaptor function contributes to cell survival under proteotoxic stress. USP18 interacts with STAT2 through isoleucine 60 (I60) to mediate negative feedback regulation of IFN signaling [24,28]. Expression of the I60N mutant abolished the protective effect observed with WT USP18 under MG132 treatment and increased proteotoxic cell death (Figures 7E, F). Notably, similar to WT USP18, the I60N mutant did not affect basal ISG expression or the MG132-induced suppression of ISGs (Figures S5C, D). Overexpression of USP18 I60N elevated its soluble pool, with a substantial fraction still partitioning into the insoluble fraction (Figure S5E), with loss of anti-apoptosis effects. These findings indicate that USP18’s cytoprotective activity is independent of its catalytic function and ubiquitination status, but critically requires its adaptor role mediated through I60 residue. We next examined whether IFNα-induced endogenous USP18 could abrogate the effects of USP18 I60N. The protective effect of IFNα against proteotoxicity was abolished following expression of the I60N mutant (Figure 7G, H). Consistent with this observation, immunoprecipitation analysis revealed markedly reduced interaction between STAT2 and USP18 I60N compared with WT USP18 (Figure 7I). Moreover, silencing STAT2 significantly impaired the cytoprotective effect of USP18 (Figures 7J, K). However, overexpression of STAT2 along with silencing of USP18 did not rescue from proteotoxic death (Figure 7L, M). Notably, overexpression of STAT2 alone under proteotoxic stress conditions failed to confer protection (Figure 7L, M), likely due to the marked downregulation of USP18 following proteasomal inhibition. In contrast, USP18 overexpression retained its protective effect under proteotoxic conditions, presumably because sufficient STAT2 protein levels were maintained despite proteasomal inhibition. We propose that this phenomenon arises from the differential sensitivity of USP18 and STAT2 to proteasomal inhibition, with USP18 being substantially more susceptible to downregulation than STAT2 (Figure 2B). Collectively, these data establish that USP18 confers resistance to proteotoxic stress through a non-enzymatic scaffolding function that is dependent on its interaction with STAT2.

## DISCUSSION

This study identifies a previously unrecognized link between proteotoxic stress and the IFN-I signaling pathway, wherein a subset of ISGs is suppressed through coordinated transcriptional and post-transcriptional mechanisms. Transcriptional repression arises from IRF9 nuclear exclusion, whereas post-transcriptional regulation is reflected by ISG partitioning into insoluble aggregates. Among these, the ISG USP18, via its scaffolding function, is essential for limiting proteotoxicity. Importantly, depletion of ISGs—especially USP18—can be reversed by exogenous IFNα, thereby preventing proteotoxic hepatocellular death (Fig 8).

**Figure 8.**
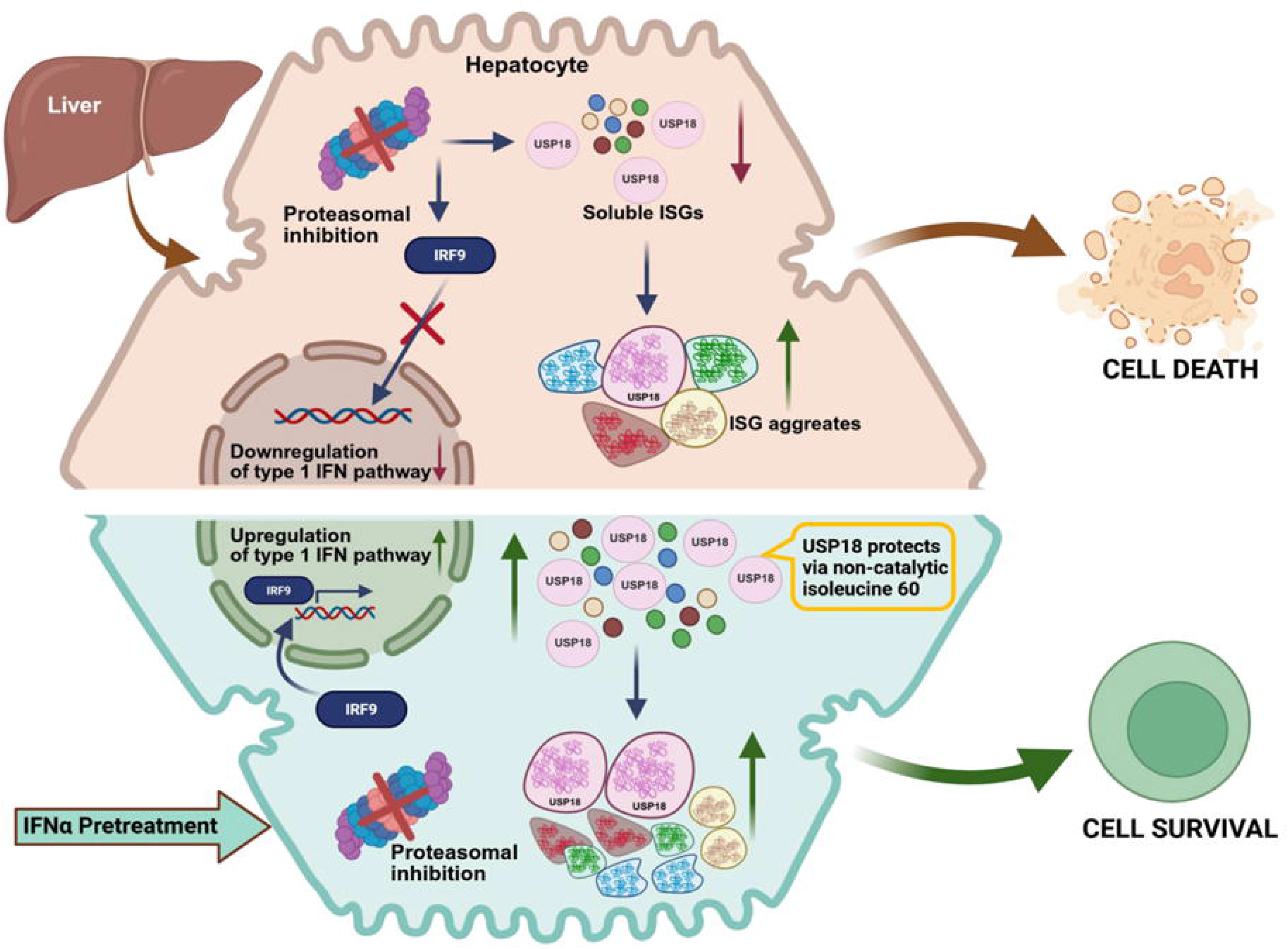
Schematic for USP18-mediated suppression of proteotoxicity.

The reciprocal regulation of IFN-I signaling and the proteasome is complex and context dependent. For instance, IFNα treatment combined with the proteasomal inhibitor bortezomib enhances cancer cell death [29–31]. Moreover, both genetic defects and chemical inhibition of the proteasome or immunoproteasome increase IFN-I production in immune cells and fibroblasts [32–33]. Plasmacytoid dendritic cells (pDCs), the primary circulatory IFN-I producers, are uniquely sensitive to bortezomib, proteasomal inhibition in pDC attenuates XBP1 splicing, disrupts ER homeostasis, impairs TLR9 trafficking, and consequently diminishes IFN-α and IL-6 production. Consistent with our data, these findings suggest that proteasome inhibitors may ameliorate inflammatory disorders such as lupus and psoriasis [34]. Additionally, IFN-α-2b protects against diet-induced obesity and attenuates MSALD by modulating SREBP-2 activity [35]. Together, these observations highlight that, beyond its canonical pro-inflammatory role, IFNα exerts non-canonical and tissue-specific functions. We further demonstrate that although IFNα has previously been reported to induce immunoproteasome complex formation, particularly during viral infections [23], we did not observe a corresponding increase in proteasomal activity in our experimental conditions. Thus, variation in cell-type susceptibility to proteasomal inhibition may critically influence downstream biological outcomes.

Proteasomal inhibition caused a marked enrichment of IFN-I signaling components and ISG proteins in the insoluble fraction. However, neither the catalytic mutant, the polyubiquitylation-deficient mutant, nor the scaffolding-defective mutant of USP18 altered its partitioning into insoluble aggregates. Instead, our findings strongly indicate that the availability of soluble USP18 is a key determinant of its cytoprotective activity under proteotoxic stress. The structural basis for ISG partitioning likely stems from IDRs, similar to those found in aggresomal proteins such as p62. IDR-containing proteins adopt expanded conformational ensembles with high structural flexibility and are particularly prone to aggregation when proteostasis is compromised. The absence of rigid tertiary structure exposes hydrophobic residues that favor intermolecular interactions and aggregation [36]. IDRs are also known to nucleate aggresome formation through liquid–liquid phase separation (LLPS) [37–38]. Consistent with an LLPS-mediated mechanism, IFN-I signaling proteins progressively regained solubility following removal of the proteasomal inhibitor, indicating that their aggregation is reversible rather than terminal.

In addition to USP18, ISG15 also mitigates proteotoxic stress; however, its role becomes dispensable when USP18 is sufficiently abundant. This observation aligns with clinical findings in individuals with inherited ISG15 deficiency, who do not exhibit heightened susceptibility to viral infection but instead display exaggerated IFN-α/β responses and autoinflammatory phenotypes reminiscent of Mendelian interferonopathies. Mechanistically, ISG15 stabilizes USP18, and its absence leads to proteasome-mediated USP18 degradation, thereby diminishing this central negative regulator of IFN-I signaling [39]. Notably, this ISG15-dependent stabilization of USP18 is species-restricted and occurs in humans but not in mice [40].

The role of USP18 in cell death regulation is highly context dependent, functioning as either pro-apoptotic or anti-apoptotic depending on the cell type and physiological state [41–44]. USP18 can promote cell survival by deubiquitinating key substrates—such as Notch1 in pancreatic cancer, SOX9 in glioblastoma, and Snail1 in colorectal cancer—preventing their proteasomal degradation and sustaining oncogenic programs [45–47]. Conversely, in lung tissue, USP18 regulates lipid and fatty acid metabolism by stabilizing ATGL and UCP1 through deISGylation, with elevated levels of these proteins correlating with tumor proliferation and poor patient prognosis [48]. Beyond cancer, USP18 also modulates metabolic disease: hepatocyte-specific overexpression of USP18 protects high-fat-diet-fed mice from MASLD by deubiquitinating TAK1 and consequently reducing JNK/NF-κB activation, inflammation, and insulin resistance [49]. In parallel, the non-enzymatic scaffolding function of USP18—particularly its role as a negative regulator of IFN-I signaling—substantially contributes to its diverse biological actions. Consistent with this, the USP18 I60N mutant exhibits defective scaffolding, thereby enhancing IFNα signaling and sensitizing cancer cells to IFN-mediated cytotoxicity [24,28]. Our observations similarly support that USP18’s scaffolding function, independent of its catalytic activity, is critical for protection against proteotoxic cell death. Importantly, human USP18 displays markedly lower enzymatic activity compared to murine USP18, suggesting that scaffolding may dominate its biological function in humans [24]. Collectively, these findings underscore the multifaceted mechanisms through which USP18 exerts context-specific physiological effects.

In summary, this study establishes a previously unrecognized connection between hepatic proteasomal inhibition and the IFN-I signaling axis, with USP18 emerging as the key molecular node integrating these pathways. The cytoprotective effect of USP18 requires its interaction with STAT2, highlighting the centrality of its scaffolding role. Although the downstream effectors recruited by USP18 during proteotoxic stress remain to be elucidated, our findings identify USP18 as a promising molecular target for mitigating proteotoxicity. Moreover, our data suggest that therapeutic augmentation of IFN-I signaling may offer a viable strategy for enhancing cellular resilience to proteotoxic stress.

## STATEMENTS & DECLARATIONS

## Author Contributions

AS performed most of the cell-based and animal experiments and responsible for acquisition of data. SC conducted IP and phase separation experiments, and analyzed the data. PC conceptualized and supervised the project and acquired research funds. AS and PC analyzed the data and wrote the paper.

## Data Availability

The datasets analysed during the current study are available in the Gene Expression Omnibus repository, file no. GSE164508.

## Ethics approval

All experimental procedures were approved by the Institutional Animal Ethics Committee of CSIR-IICB and complied with CPCSEA guidelines, Government of India.

## Acknowledgements

We thank Sounak Bhattacharya and Rabin Pramanik for assisting with confocal microscopy and biochemical experiments, respectively. We also thank Central Instrumentation Facility (CIF) at CSIR-IICB for support with various equipments.

## METHODS

### Cell culture

Hepatocellular carcinoma cell line HepG2 cells were purchased from ATCC [ #HB-8065] and grown in Minimum Essential Media (MEM) [Himedia, Mumbai, India, #AL047S] with10% FBS [GibcoThermo Fisher Scientific, MA, USA, #10270106] and 1% antibiotic solution [HiMedia, Mumbai, India, #A018] in a humidified incubator with 5% CO_2_ at 37°C. Human embryonic kidney cell line HEK293A was cultured in Dulbecco’s Modified Eagle medium (DMEM) [Himedia, Mumbai, India, #AL007S] with 10% FBS, 1% Antibiotic-Antimycotic, and 1% NEAA [Himedia, Mumbai, India, # ACL006] at 37°C in 5% CO_2_.

### Purification of Recombinant USP18 Adenovirus

Recombinant adenovirus carrying mouse USP18 [abm, Canada, #492950540200] and eGFP transgene were amplified and propagated in HEK293A cells, and infected cell pellets were processed using PureVirus™ Adenovirus Purification Kit [Cell Biolabs, CA, USA, #VPK-5112]. The final eluate was passed through a microspin column, supplemented with 10% glycerol, and stored at −80°C. Viral titers were determined spectrophotometrically at 260 nm, using the formula: pfu/ml = 1.1 × 10¹² × OD₂₆₀ × dilution factor.

### Animal experiment

Wild-type male C57BL/6 mice (6–8 weeks old) were housed in individually ventilated cages under standard conditions (12 h light/dark cycle, 22 ± 1 °C) with ad libitum access to food and water. Mice were randomly assigned to different experimental groups. No blinding was applied during the experimental procedures, and no predefined exclusion criteria were used. All experimental procedures were approved by the Institutional Animal Ethics Committee of CSIR-IICB and complied with CPCSEA guidelines, Government of India. Mice were intraperitoneally injected with Bortezomib [Merck, MA, USA, #5043140001] or IFNα [Biolegend, California, USA, #752804], and liver and blood were collected. In order to overexpress USP18 in mouse liver, mice were injected intravenously via tail vein with recombinant adenovirus expressing eGFP and USP18 (∼ 2X10^11^ pfu).

### Antibodies

Following antibodies were purchased from Cell Signaling Technologies (CST, MA, USA) and used for immunoblotting: Stat1 (D1K9Y) [#14994], Stat2 (D9J7L) [#72604], pSTAT1 (Tyr701) (58D6) [#9167], pSTAT2 (Tyr690) (D3P2P) [#88410], IRF9 (D2T8M) [#76684], IRF9 (D9I5H) [#28845], USP18 (D4E7) [#4813], USP18 (E9K4X) [#53229], ISG15 [#2743], Ubiquitin (P37) [#58395], DYKDDDDK Tag (D6W5B) [#14793], HA-Tag (C29F4) [#3724], SQSTM1/p62 (D5L7G) [#88588],Histone H3 (D1H2) [#4499], GAPDH mAb [#AC002], Cleaved Caspase-3 (Asp175) Antibody [#9661S], Cleaved PARP (Asp214) (D6X6X) [#94885]. Mouse β-Actin was purchased from Sigma MO, USA [#A5316], and Anti-rabbit secondary antibody HRP tag [#31460], Anti-mouse secondary antibody HRP tag [#31430] were purchased from Invitrogen [MA, USA].

### Transfection

HepG2 cells were forward transfected with required amount of plasmid DNA -Flag USP18 [Genescript, NJ, USA #OHu15965], HA ISG15 [Addgene, MA, USA, #80404], HA Ubiquitin (Ub) [Addgene, MA, USA, # 18712] using Lipofectamine 2000 (Invitrogen, MA, USA#11668019) reagent in Opti-MEM media [Gibco, Invitrogen, MA, USA, #31985070] at a ratio of 1:3 [1μg plasmid DNA: 3μl Lipofectamine 2000]. siRNA was reverse-transfected using Lipofectamine RNAiMAX reagent [Invitrogen, MA, USA, #13778150]. Smart Pool siRNA against human STAT1 [#L-003543], STAT2 [#L-012064], USP18 [L-004236], ISG15 [L-004235], and IRF9 [L-020858] were purchased from Dharmancon Horizon Discovery [CO, USA].

### Protein estimation and extraction

HepG2, HEK293A, and liver were lysed or homogenized in buffer (100 mM NaCl, 50 mM Tris-HCl, 1 mM EGTA, 1 mM EDTA, 1% Triton X-100, protease [Merck Millipore, MA, USA, #539134] and phosphatase inhibitors [Roche, Basel, Switzerland, #4906837001] and centrifuged at 20,000g for 20 min at 4°C to obtain detergent soluble proteins. The pellet was washed, resuspended in lysis buffer containing 1% SDS, heated at 95°C for 10 min, and centrifuged to obtain the detergent insoluble fraction [49]. Protein concentrations were determined using BCA assays [Thermo Fisher, MA, USA, #23235].

### Inhibitors

MG132 [Sigma. MO, USA, #M7449] was used at a standard dose of 5μM across experiments [7,13]. Z-VAD(OMe)-FMK [Cayman, MI, USA, #14463] was used at a final dose of 50µM. Bafilomycin A1 [ Sigma. MO, USA, #B1793] was used at a final dose of 50 nM. 5% 1,6 Hexanediol [ Sigma, MO, USA, #240117] was used for 2 min [50].

### Western blotting

Proteins were separated on 8–15% SDS-PAGE gels and transferred to 0.45 μm PVDF membranes [Merck Millipore, MA, USA, #IPVH00010]. Membranes were blocked with 5% skim milk [Himedia, Mumbai, India, #GRM1254] in tris-buffered saline with tween-20 (TBST), and incubated overnight at 4°C with primary antibodies (1:1000 in 5% BSA + 0.04% sodium azide). After washing, membranes were incubated with HRP-conjugated secondary antibodies at a dilution of 1:2000 in 5% milk for 1 h, washed, and images were developed using Clarity™ ECL [BioRad, CA, USA] in iBright 1500 system [Invitrogen Thermo Fisher Scientific, MA, USA]. At least n=3 independent experiments were performed for all Western blot assays.

### Immunoprecipitation

For the overexpressed proteins, HepG2 cells were transfected with plasmids, and after 48 h, cells were treated with 5μM MG132 [Sigma, CO, USA, #M7449] for 6h. For immunoprecipitation, 200 μg protein was incubated overnight at 4°C with 20μl Protein A/G magnetic beads [Merck Millipore Milliopore, MA, USA, #LSKMAGA02] and 2 μg of flag-tag antibody. Beads were washed and eluted with 2× loading buffer, and samples were analyzed by immunoblotting. For endogenous USP18, 1 mg of HepG2 cell lysates were immunoprecipitated using anti-USP18 [CST, MA, USA, #4813] antibody at a dilution of 1:50 and blotted with anti-ubiquitin antibody.

### RNA isolation and cDNA synthesis

Cultured cells and tissues were lysed or homogenized in TRIzol [Invitrogen, MA, USA, #15596018], centrifuged, and the supernatant was treated with chloroform. After phase separation, RNA from the aqueous phase was precipitated with isopropanol, washed with 70% ethanol, air-dried, and dissolved in DEPC-treated water. For cDNA synthesis, 1 μg RNA was reverse transcribed using iScript Supermix [Bio-Rad, CA, USA, #1708841].

### Quantitative PCR (qPCR)

Synthesized cDNA was diluted with DEPC-treated water (1:1). Gene-specific primers were designed using Primer3, synthesised by Integrated DNA Technologies (IDT, Lowa, USA). qPCR reaction was conducted using 2× SYBR Green Master Mix [Bio-Rad, CA, USA, #1725124] for 40 cycles on a Roche Light Cycler 96. The primers used are included in Supplementary Table 1.

### Immunofluorescence

Following treatment, cells were fixed in 4% paraformaldehyde, permeabilized, and blocked with 5% serum in which the secondary antibody was raised, and 0.3% Triton X-100. Cells were incubated overnight at 4°C with primary antibodies followed by fluorescence conjugated secondary antibodies. Nuclei were counterstained with 25 μg/ml Hoechst 33342 [Invitrogen, MA, USA, #H3570]. Images were captured using a Leica TCS SP8 STED confocal microscope. Antibodies used are- USP18 pAb [ABclonal, MA, USA, #A16739], CST (MA, USA) antibodies IRF9 (D2T8M) [#76684], DYKDDDDK Tag (9A3) [#8146], SQSTM1/p62 (D5L7G) [#88588], Cleaved caspase-3 (Asp175) (D3E9) mAb (Alexa Fluor® 647 Conjugate) [#9602S]. Anti-mouse secondary antibody Alexa Fluor 647 [Invitrogen, MA, USA, #21236], Anti-rabbit secondary antibody Alexa Fluor 647 [Invitrogen, MA, USA, #A21245], Anti-rabbit secondary antibody Alexa Fluor 488 [Invitrogen, MA, USA, #A11034].

### Immunohistochemistry

Paraffin sections were heated at 85°C, deparaffinized, rehydrated, and antigen retrieval was performed in sodium citrate buffer (10 mM, 0.05% Tween-20). Slides were blocked in 1.5% serumderived from the host species in which the secondary antibody is raised. Slides were incubated overnight at 4°C with primary antibodies. For detection, either VECTASTAIN ABC Kit or ImPRESS Duet Kit [Vector Labs, CA, USA, # PK-6100] was used, depending upon the species in which the primary antibody was raised, followed by addition of 3,3′-Diaminobenzidine(DAB) [Vector Labs, CA, USA, #SK-4100]substrate. Nuclei were counterstained with hematoxylin [Vector Labs, # H-3404], mounted using Vectamount AQ [Vector Labs, CA, USA, #H-5501] and images were acquired under an EVOS XL Core microscope [Thermo Fisher, MA, USA].

### AST and ALT

Blood was collected from the sacrificed C57BL/6 mice by cardiac puncture. The serum was collected, and alanine aminotransferase (ALT) and aspartate aminotransferase (AST) were analyzed using the manufacturer’s protocol (Randox, UK, #AL2780, #AS2782).

### Live Dead cellular staining assay

Cells were stained with 2 μM Calcein AM for live cells (green) and 4 μM Ethidium homodimer-1 for dead cells (red) in PBS for 30 min using the LIVE/DEAD™ Viability/Cytotoxicity Kit [Invitrogen, MA, USA, #L3224]. Fluorescence was imaged using a Leica TCS SP8 STED confocal microscope with excitation/emission filters of 494/517 nm and 528/617 nm.

### Quantification of Apoptosis and Viability

HepG2 cells were reverse-transfected with 5 pmol of siRNAs and were seeded in an opaque 96-well plate with transparent bottom (20,000 cells/well; Thermo Fisher, #165305). Cell viability and apoptosis were measured sequentially using the ApoLive-Glo™ Multiplex Assay [Promega, Wisconsin, USA, #G6410]—fluorescence (Ex 400/Em 505 nm) measured cell viability, and luminescence measured caspase-3/7 activity. The caspase-to-viability ratio reflected normalized apoptotic response.

### Nuclear protein extraction

Nuclear and cytoplasmic fractions were prepared from fully confluent HepG2 cells using a Nuclear Extraction Kit [Cayman, MI, USA, #10009277] as per the manufacturer’s protocol. Protein was measured by BCA assay [Thermo Fisher, MA, USA, #23235] for subsequent immunoblot analysis.

### Protein droplet assay

Flag-tagged USP18 was transfected in HEK293A cells, and cells were treated with 5 µM MG132 for 12h. Proteins were extracted, and 4 mg lysate from control and treated cells was immunoprecipitated using 4 µg anti-flag antibody [Sigma, MO, USA, #F1804] or IgG control. USP18 was eluted with 500 µg/ml Flag peptide [Sigma, MO, USA, #F3290]. For droplet visualization, 96-well plates were pre-coated overnight with 30 mg/ml fatty acid-free BSA [22] [MP Biomedical, CA, USA, #152401]. Purified USP18 from control and MG132-treated cells was incubated along with pure mouse IgG in another well for 2 h at 37°C, and droplet formation was imaged using the Evos XL Core microscope [Thermo Fisher, CA, USA].

### Actinomycin D assay

HepG2 cells were treated with 5µg/ml dose of Actinomycin D [Sigma, MO, USA, #114666] dissolved in DMSO for 2h, 4h, 8h and 12h. Another group of cells were treated with 10µM of MG132 along with the above dose and time of Actinomycin D.

### Caspase 3/7 activity assay

Cellular Caspase-3/7 activity was detected using CellEvent Caspase-3/7 Green Detection Reagent [Invitrogen, MA, USA, #C10423] as per manufacturer’s protocol. Nuclei were stained with 25μg/ml of Hoechst 33342. Fluorescent images were captured using a Leica TCS SP8 STED confocal microscope with Ex 502/Em 530 nm.

### Site-directed mutagenesis

The cysteine residues at position 64 in USP18’s active site were mutated to alanine to generate a catalytically inactive USP18. Additionally, lysine residues were mutated to generate USP18 lysine mutants. Mutagenesis was performed using 2X Q5® Hot Start High-Fidelity DNA Polymerase [NEB, MA, USA, #M0494S] with 50 ng plasmid DNA, 500 nM primers, and 1× master mix. The primer sequences are available in Supplementary Table 2.

### Transcriptomics data analysis

Hepatic transcriptomic data from vehicle and bortezomib-treated mice were analyzed using DESeq2 file (GSE164508) [7]. Differentially expressed genes (log2FC, padj) were subjected to GO and KEGG enrichment using *clusterProfiler* and *org.Mm.eg.db* in R Studio 2025.05.1, focusing on interferon-related pathways (padj< 0.05). Pathway directionality was determined from up- and downregulated enrichments, and visualized as diverging barplots in *ggplot2*. Volcano plots were generated in R using log₂ fold change versus −log₁₀(adjusted p-value). Genes with adjusted p-value< 0.05 and |log₂ fold change| ≥ 1 were considered significant. ISGs were identified using a curated gene list from MSigDB by including the genes of hallmark interferon alpha response and response to IFN-α gene sets. ISG heatmaps were plotted in GraphPad Prism using Z-scores.

### Statistics

All data are represented as mean ± SD, and GraphPad Prism (8.0.1) was used for all analysis. Unpaired two-tailed *t*-test with Welch correction and two-way ANOVA were performed, with significance defined as p ≤ 0.05. For microscopy analysis, the ImageJ platform was used to calculate mean fluorescence intensity (MFI) and particle number. Densitometry of western blots was analyzed by ImageJ. Relative abundance of soluble and insoluble proteins was determined by densitometric analysis, considering the soluble fraction at the basal state as 1. Sample size and variance were not predetermined using statistical methods.

