## SUPPLEMENTARY FIG AND TABLES for "USP18-STAT2 axis enhances hepatic resilience under proteotoxic stress"

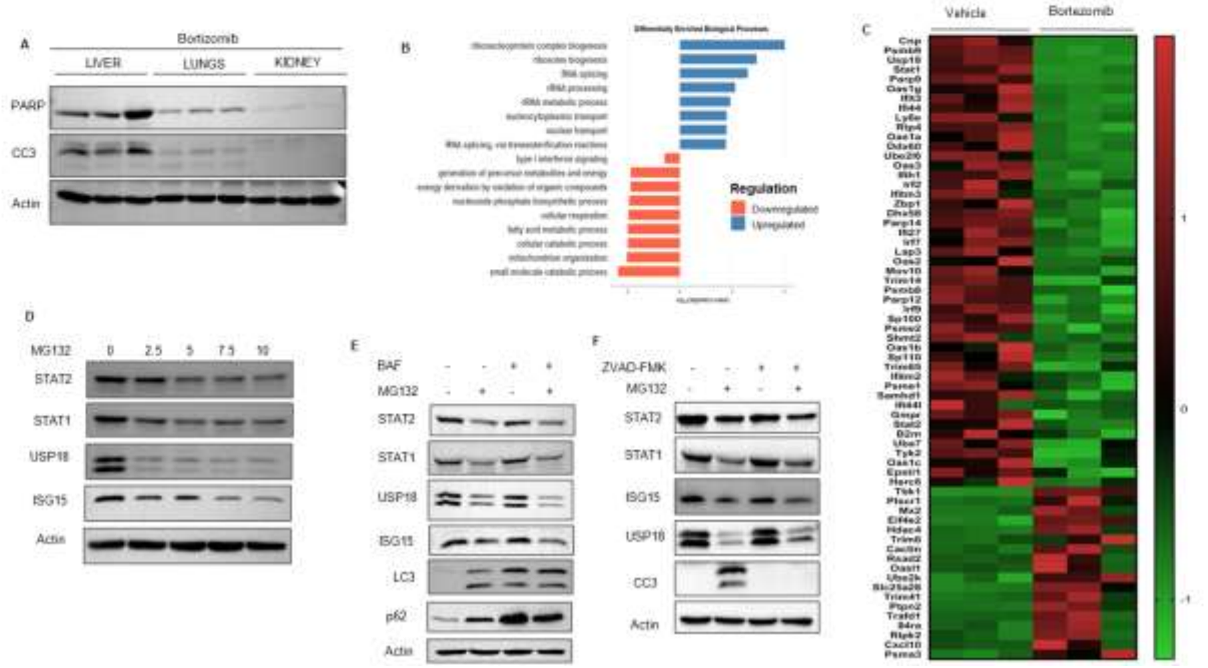


**Figure S1. Type 1 interferon signaling pathway and proteasome.**

(A) Western blot for apoptosis detection in the indicated organs by IP injection of 5mg/kg body weight of bortezomib in C57BL/6 mice (n=3 mice). Each lane represents tissue from one individual mouse. (B) Diverging bar plot showing significantly enriched Gene Ontology biological processes based on $-\log_{10}$ values (padj $< 0.05$). (C) Heat Map showing differentially regulated ISGs in liver on bortezomib treatment in C57BL/6 mice for 16h (padj $< 0.05$). (D) Western blot analysis of lysate obtained from HepG2 cells treated with the indicated dosage of MG132 for 16h. **(**E) HepG2 cell treated with 50 nM bafilomycin A for 4h followed by 5µM of MG132 treatment for 16h, and analyzed by immunoblotting. (F) HepG2 cell treated with 50µM of Z-VAD(OMe)-FMK for 4h, followed by 5µM of MG132 treatment for 16h and analyzed by immunoblotting.

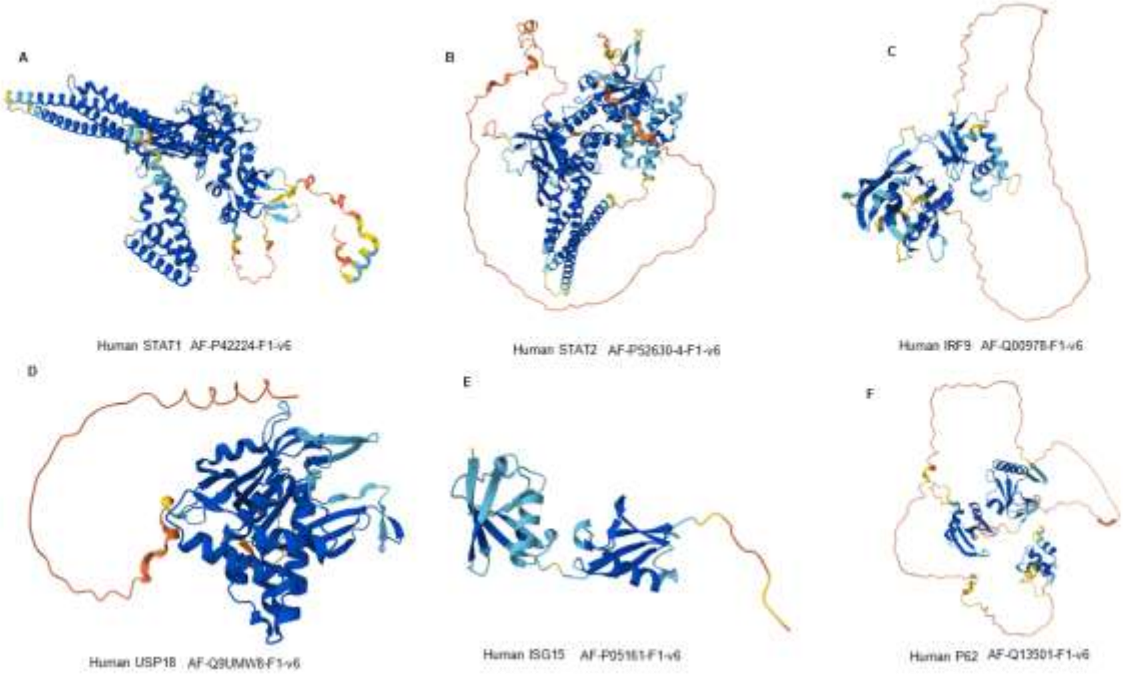


**Figure S2. Intrinsically disordered regions of ISGs.** Structure generated from Alphafold.

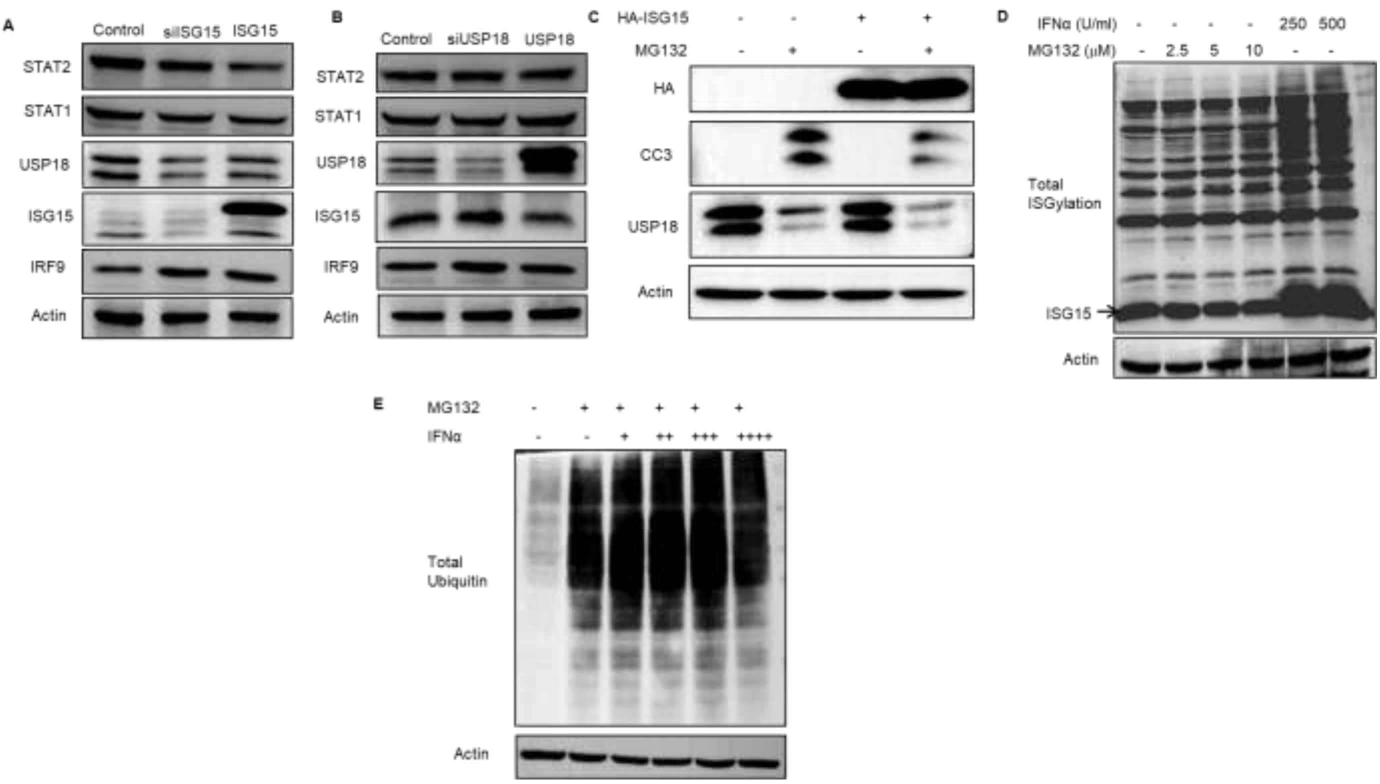


**Figure S3. Impact of USP18 and ISG15 on the ISGs expression.**

(A) Western blot analysis of lysate from HepG2 cells transfected with siISG15 and ISG15 for 48h. (B) Western blot analysis of lysate from HepG2 cells transfected with siUSP18 and USP18 for 48h. (C) Effect of ISG15 overexpression in proteotoxicity mediated apoptosis. (D) Western blot analysis in HepG2 cells treated with 5µM of MG132 and 250, 500 IU/ml of IFNα for 16h to check the total ISGylation. (E) Western blot analysis in HepG2 cells pre-treated with 100, 250,500 and 1000 IU/ml concentration of IFNα for 4h, followed by 12 h of 5µM of MG132.

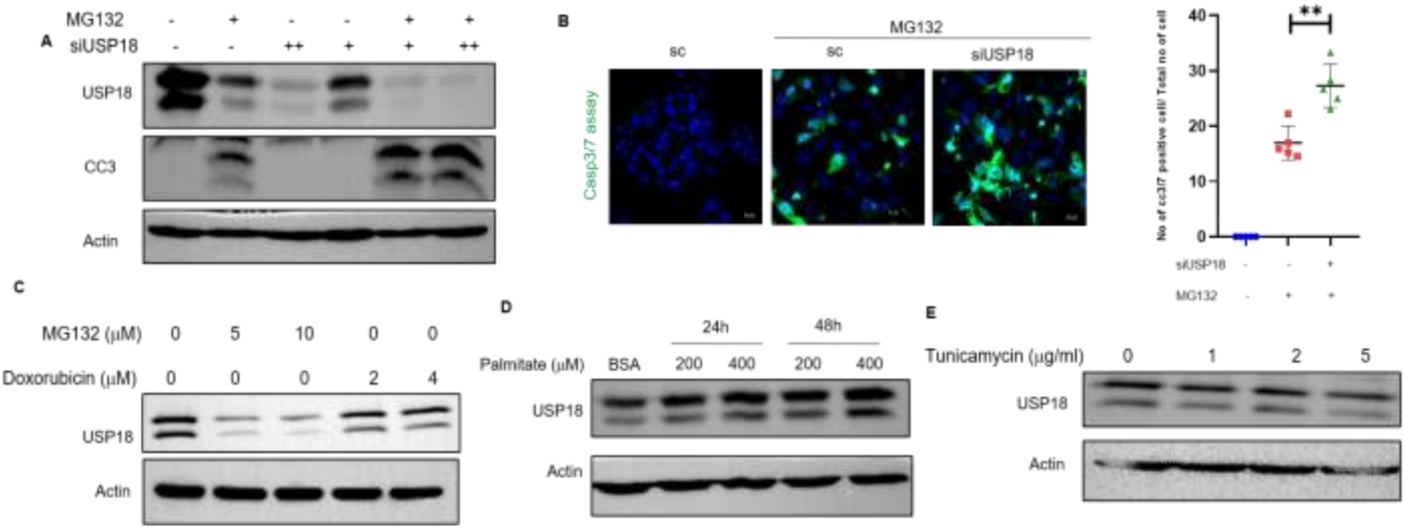


**Figure S4. Specificity of USP18 to proteotoxic cell death.**

(A) HepG2 cells were transfected with 50 and 100 pmole of siUSP18. 56h post transfection 5μM of MG132 treatment was given for 16 h. Cell lysates were analysed by immnoblotting. (B) Caspase3/7 activity in cells transfected with 50 pmole of siUSP18 for 56h followed by 16h of 5μM of MG132 treatment. Values represent mean ± SD (5 fields/group). Statistical significance: **p< 0.01. Scale bar = 10 μm. (C) USP18 in genotoxic stress. HepG2 cells were treated with 10μM of doxorubicin for 24h**.** **(**D) Role of USP18 in ER stress. HepG2 cells were treated with 2 μg/ml and 5μg/ml of tunicamycin for 16h. (E**)** USP18 expression in lipotoxicity**.** HepG2 cells were treated with 200μM and 400μM of palmitic acid for 24h and 48h. BSA served as vehicle control.

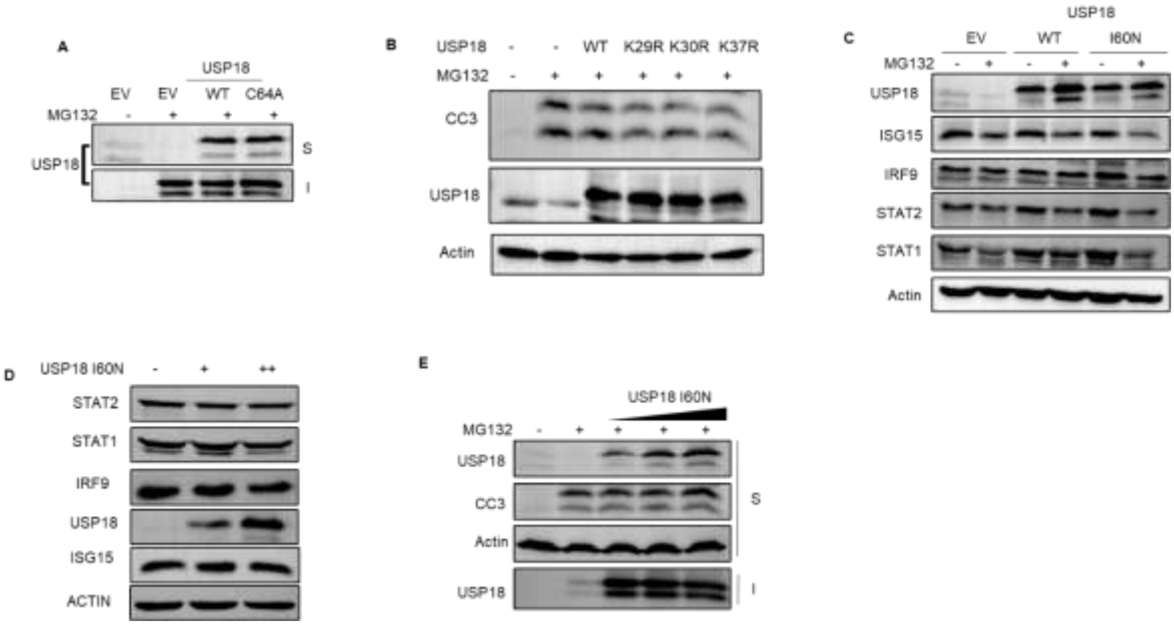


**Figure S5. Role of USP18 lysine and I60N mutants in proteotoxicity.**

(A) Western blot for soluble and insoluble fractions of USP18 catalytic mutant. (B) Western blot showing the effect of three lysine mutants in proteotoxic death. (C) Effect of WT USP18 and I60N mutant on ISG expression under proteotoxic stress. (D) Dose-dependent effect of mutant USP18 I60N on other ISGs. (E) Dose-dependent effect of I60N mutant in proteotoxic stress-mediated apoptosis. S-detergent soluble fraction, I-detergent insoluble fraction.

**Table 1.** List primers used for qPCR

| GENE | Forward Primer (5'-3') | Reverse Primer (5'-3') |
|---|---|---|
| **18S (Human)** | GTAACCCGTTGAACCCCATT | CCATCCAATCGGTAGTAGCG |
| **STAT1 (Human)** | TGTGAAGTTGAGAGATGTGA | CTTCAGTAACGATGAGAGGA |
| **STAT2(Human)** | AACATTCTGACTTCAAACCA | GTATATTTGACCGTGAAGCT |
| **IRF9 (Human)** | TGAATTTCTGGGAAGAGAGC | CAGGAAGAACAAAGAGGTGA |
| **USP18 (Human)** | GCTCCTGAGGCAAATCTGTC | CTGGGCTGCTCTCTCTTCAT |
| **RTP4 (Human)** | TCTCCAGAAATGCCAGTAAT | AATTCCCAGTCTTTAGGTGG |
| **ISG15(Human)** | GCAGATCACCCAGAAGAT | GCCCTTGTTATTCCTCAC |
| **18S (Mouse)** | GTTGGTTTTCGGAACTGAGG | TCGTTTATGGTCGGAACTACG |
| **STAT1 (Mouse)** | TAATTTCTTGTTGCAGCACA | CACATGACTTGATCCTTCAC |
| **STAT2(Mouse)** | CAGAGACAGGGCTTAATTTG | TGGAGAGTTGGTTCATGTTA |
| **IRF9 (Mouse)** | CAAGAGTTCCGAATTTGAGG | GACACACAACTGATACTTCG |
| **USP18 (Mouse)** | AAGGACCAGATCACGGACAC | CACATGTCGGAGCTTGCTAA |
| **RTP4 (Mouse)** | GAGATACTATGGACACAGGA | TTGGTAATGGAGATGGAGAT |
| **ISG 15 (Mouse)** | GTGACTAACTCCATGACG | CAGAAAGACCTCATAGATGT |

**Table 2.** List of primers used for site-directed mutagenesis (SDM).

| GENE | Forward Primer (5'-3') R | Reverse Primer (5'-3') |
|---|---|---|
| **USP18 SDM C64A** | CATTGGACAGACCGCCTGCCTTAACTCC | GGAGTTAAGGCAGGCGGTCTGTCCAATG |
| **USP18 SDM I60N** | TGGTTTACACAACAACGGACAGACCTGCT | AGCAGGTCTGTCCGTTGTTGTGTAAACCA |
| **USP18 SDM K29R** | GCAGATCTTGAAGAAAGGAAGGAAGAAGACAGC | GCTGTCTTCTTCCTTCCTTTCTTCAAGATCTGC |
| **USP18 SDM K30R** | GATCTTGAAGAAAAGAGGGAAGAAGACAGCAAC | GTTGCTGTCTTCTTCCCTCTTTTCTTCAAGATC |
| **USP18 SDM K37R** | ACAGCAACATGAGGAGAGAGCAGCC | GGCTGCTCTCTCCTCATGTTGCTGT |
